# Statistical Approaches to Decreasing the Discrepancy of Non-detects in qPCR Data

**DOI:** 10.1101/231621

**Authors:** Valeriia Sherina, Helene R. McMurray, Winslow Powers, Hartmut Land, Tanzy M.T. Love, Matthew N. McCall

## Abstract

Quantitative real-time PCR (qPCR) is one of the most widely used methods to measure gene expression. Despite extensive research in qPCR laboratory protocols, normalization, and statistical analysis, little attention has been given to qPCR non-detects – those reactions failing to produce a minimum amount of signal. While most current software replaces these non-detects with a value representing the limit of detection, recent work suggests that this introduces substantial bias in estimation of both absolute and differential expression. Recently developed single imputation procedures, while better than previously used methods, underestimate residual variance, which can lead to anti-conservative inference. We propose to treat non-detects as non-random missing data, model the missing data mechanism, and use this model to impute missing values or obtain direct estimates of relevant model parameters. To account for the uncertainty inherent in the imputation, we propose a multiple imputation procedure, which provides a set of plausible values for each non-detect. In the proposed modeling framework, there are three sources of uncertainty: parameter estimation, the missing data mechanism, and measurement error. All three sources of variability are incorporated in the multiple imputation and direct estimation algorithms. We demonstrate the applicability of these methods on three real qPCR data sets and perform an extensive simulation study to assess model sensitivity to misspecification of the missing data mechanism, to the number of replicates within the sample, and to the overall size of the data set. The proposed methods result in unbiased estimates of the model parameters; therefore, these approaches may be beneficial when estimating both absolute and differential gene expression. The developed methods are implemented in the R/Bioconductor package nondetects. The statistical methods introduced here reduce discrepancies in gene expression values derived from qPCR experiments, providing more confidence in generating scientific hypotheses and performing downstream analysis.

## 1 Introduction

Polymerase chain reaction (PCR), developed over 30 years ago by Dr. Kary Mullis, uses short-length oligonucleotide *primers* to initiate and direct synthesis of new DNA copies using DNA polymerase plus single-stranded DNA as a template [21]. Oligonucleotides complementary to each of the two possible sequences relating to the sense and anti-sense strands of the target DNA are included in the reaction, allowing both strands to be amplified simultaneously. These new DNA copies are added to the pool of DNA templates and the process is repeated multiple times, so that amplification occurs by chain reaction [2].

The use of fluorescent tags allows one to quantify the amount of DNA present at each PCR cycle; this is referred to as *quantitative PCR* or *qPCR*. Unlike other genomic technologies, qPCR allows one to measure the rate of amplification across PCR cycles [29]. Analysis of the resulting amplification data across PCR cycles can be used to estimate a quantity proportional to the initial amount of DNA in a sample, referred to as the quantification cycle (Cq). The most widely used Cq value is the threshold cycle (Ct), the cycle at which expression of a target gene first exceeds a predetermined fluorescence threshold level. This quantity is inversely proportional to the number of target molecules in the initial pool; therefore, a higher Ct value implies there was lower expression of the target in a sample and more PCR cycles were required to exceed the fluorescence threshold. Ct values are either related to a known set of copy number standards or a control gene (absolute quantification) [20, 23] or to the Ct value of the same target in another sample (relative quantification) [24].

In addition to quantifying DNA, qPCR can be used to measure gene expression at the level of mRNA by first generating a complementary DNA (cDNA) via reverse transcription of the pool of mRNA and then using the cDNA for target amplification by PCR. qPCR remains the gold standard for measuring gene expression below genome-scale [16]. qPCR is widely used in gene therapy distribution and expression assays [30], drug response analysis [1, 13], mutation analysis [11, 28], single nucleotide polymorphism (SNP) analysis [14, 10, 25], expression system development [22], detection and quantification of pathogens [12, 26], GMO detection in food [3, 19], human and veterinary diagnostics [27, 17], as well as in fetal medicine [5].

### 1.1 Statistical challenges in qPCR data analysis

An important issue in qPCR experiments that has been largely ignored is the presence of non-detects, those reactions failing to pass the given quantification threshold and therefore lacking a Ct value. There are several opinions about the nature of non-detects and how to deal with their presence. Integromics RealTime StatMiner divides non-detects into *absent values*, those samples for which reactions did not occur, and *undetermined values*, those reactions that failed to pass a predetermined quantification threshold. Absent values are set to the median value of the replicates detected in the set of experiments, and undetermined values are replaced by the maximum possible Ct value (typically 35 or 40) [9]. The Applied Biosystems DataAssist v3.0 software allows users to set a Maximum Allowable Ct Value, a cut-off such that any value greater than this chosen threshold will be changed to this maximum allowable value.

If we assume non-detects occur completely at random, then simply removing them would lead to unbiased and consistent estimates of the RNA expression. Alternatively, if we assume non-detects are missing at random given the expression in replicate samples, a mean imputation procedure, in which missing values are replaced by their conditional expectation, would produce unbiased and consistent estimates of average expression. However, this approach would distort the distribution of gene expression and lead to underestimation of residual variance [7].

Recent work suggests that neither of these assumptions are valid for qPCR non-detects. Previously, it has been shown that the probability of a non-detect increases as the expression of the target transcript decreases; therefore, non-detects do not occur completely at random [18]. While it is often not possible to distinguish between missing at random and missing not at random from the observed data, previous work suggests that qPCR non-detects are likely missing not at random based on specific aspects of the technology and prior analysis of a large control data set [18]. We have developed a single imputation procedure that treats non-detects as non-random missing data, models the missing mechanism as a monotone increasing function of gene expression, and uses an Expectation-Maximization (EM) algorithm to impute missing values. This approach was a significant improvement over previous approaches, but it underestimates the residual variance, leading to anti-conservative inference.

We propose to address this issue by implementing an algorithm to directly estimate the mean and variance of gene expression using Maximum Likelihood Estimation (DirEst). Additionally, we have extended the methodology from a Single Imputation (SI) to a Multiple Imputation (MI) procedure that accounts for the uncertainty in the imputed values. We have incorporated three sources of variability into our model: uncertainty in the missing mechanism, uncertainty in the parameter estimates, and measurement error.

In Section 2 we propose methods for handling non-detects in the qPCR data and introduce models for absolute and relative quantification of the gene expression. The model for relative quantification allows one to adjust for a potential batch effect. The model for the missing data mechanism is also given in this section. We emphasize that this model arises from a mechanistic understanding of the biochemical processes in the qPCR experiments. Section 3 contains assessments of the proposed method using both simulations and three real data sets. We study our procedures’ robustness to model assumptions and compare them to other existing methods.

## 2 Materials and Methods

In this manuscript we propose two new methods to handle qPCR non-detects that provide consistent estimates of the first and second moments of gene expression: MI and DirEst. MI does this by taking into account the uncertainty of the imputed data. DirEst allows one to directly estimate within replicate means and variances for each gene and sample type. This approach is applicable to the most common subsequent analyses, e.g. identification of genes that are differentially expressed between sample types, for which the within-replicate means and variances are sufficient statistics, meaning that all the information about the data log-likelihood is contained in these parameters [6]. The limitation of the DirEst approach is that individual expression values are unavailable, so clustering or coexpression analyses are not possible.

### 2.1 A statistical model for qPCR non-detects

We propose the following generative model for qPCR data in which *Y_ij_* is the measured expression value for gene *i* in sample *j*, some of which are missing (non-detects), *X_ij_* represents the fully observed expression values, and *Z_ij_* indicates whether a value is detected:

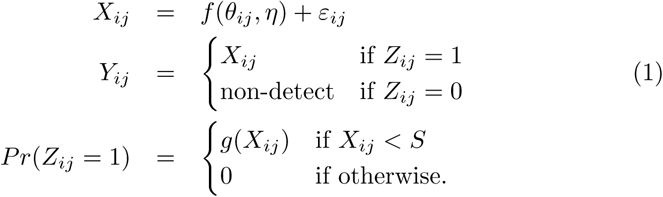

In this model, we assume that the fully observed expression values, *X_ij_* are a function of the true gene expression, *θ_ij_*, non-biological factors, *η*, and random measurement error, ε_ij_. The probability of an expressed value being detected is assumed to be a function of the expression value itself, *g*(*X_i_j*), for values below the detection limit, *S*. The following logistic regression model is a natural choice for such a relationship:

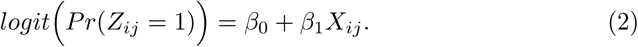

The vast majority of qPCR experiments seek to compare replicate samples from two or more sample-types. The model proposed above can be easily tailored to this type of experimental design. Specifically, we partition the samples (*j* = 1,…, *J*) into *K* sets of replicates, J_k_, with *k(j*) = *k* for *j* ∊ *J_k_*. In Equation 1, we simply replace *θ_ij_* with *θ_ik(j)_*. In the following two subsections, we describe special cases of the proposed model to estimate absolute or relative expression.

#### 2.1.1 Model for absolute quantification

Absolute quantification is used to estimate the expression of a target transcript in one or more sample-types. For *absolute* gene expression we write the proposed model as follows:

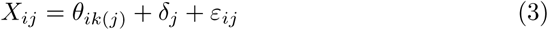

where *X_ij_* are again the completely observed gene expression values, *θ*_*ik*(*j*)_ are the true values of gene expression for gene *i* in the sample-type *k* to which sample *j* belongs, *δ_j_* represents a global shift in expression across samples, and *ε_ij_* now captures both biological and technical variability. In addition to estimating absolute expression within each sample-type, parameter estimates from this model can be used to assess relative expression between sample-types.

#### 2.1.2 Model for relative quantification

Due to the significant impact of batch effects on genomic data [15], it is increasingly common for experiments to include a matched control sample for each sample or group of samples analyzed. In this case, the parameters of interest are no longer the average expression within each sample type; rather, they are the differences in expression between the test and control samples. These control samples can be included in a model to directly adjust for batch effects. Specifically, we partition samples into 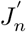 batches, with *n*(*j*) = *n* for 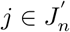, and introduce *γ_in_* into the model to capture the batch effect for gene i in samples from batch *n* as follows:

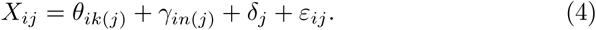

Here, the parameter of interest, *θ_ik_*(*j*), is the difference in expression between the test and control samples. Specifically, in Equation 4, we assume that control samples are denoted by *k* = 0 and *θ*_*i*0_ = 0 ∀*i*, such that:

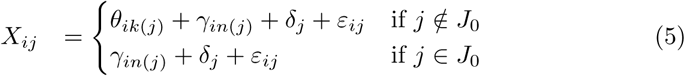

Rarely, non-detects may occur in the control samples as well as in the test samples. In this case we propose to use a two step process: first, apply the model in equation 3 for the control samples and perform SI; second, use the model in equation 4, to obtain the estimates of the non-detects for the test samples.

### 2.2 Improved estimation of the missing data mechanism

It is possible to observe perfect separation between observed and non-detected transcripts. This is a common problem in applied regression with binary predictors. Prediction of the parameter values in this case becomes unstable. [8] proposed to use Bayesian inference to obtain stable estimates of the generalized linear regression coefficients, and we adopted their approach while performing SI, MI and DirEst procedures.

### 2.3 Multiple Imputation

The central idea of MI is to replace a set of missing data points with M sets of plausible values. These sets of values are independent between the imputations, but share a correlation structure within each complete data set. This method captures the uncertainty in the imputed values. The resulting *M* complete data sets can be analyzed using standard statistical techniques, and the results can be combined and compared across the *M* datasets. Contradicting results may indicate that the imputed values are driving the inference and not the observed data.

We incorporated different sources of variability in the MI procedure: random noise, uncertainty in the linear model parameters (*θ* in Equations 3 & 4), and parameters of the logistic regression model (*β*_0_ & *β*_1_ in Equation 2), as well as each combination of these sources of variability.

#### 2.3.1 Uncertainty in linear model parameters

When the data has missing values, the estimates of the model parameters contain an additional amount of uncertainty due to the missing data. One can account for this added uncertainty by introducing additional variation in the parameter estimates. Instead of using point estimates 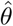 we draw *M* different 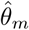 (*m* = 1,…, *M*) from the estimated distribution of 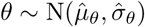.

#### 2.3.2 Uncertainty in the missing data mechanism

Similarly, one can account for the uncertainty in the missing data mechanism by introducing additional variability in the corresponding parameters estimates. To preserve the dependence between the parameters, we assume (*β*_0_,*β*_1_) are jointly MVN 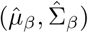. One can draw M pairs of (*β*_0_,*β*_1_) from the estimated distribution and use these estimates in the imputation procedure. This step introduces additional variability in the resulting complete data sets that reflects uncertainty in the estimated logistic regression model parameters.

#### 2.3.3 Biological variability and measurement error

Suppose the model parameters are known and one is interested in applying MI to obtain estimated differential gene expressions. In this case, given all the model parameters, the imputed values will be identical and equal to the conditional expectation of the missing data point without additional variability. Such a result is undesirable, as it will lead to artificially small variance estimates. To better estimate the uncertainty of the missing value itself under known true parameters of the modeling framework, we must include biological variability and measurement error in the MI procedure. We assume that these sources of variability together are normally distributed with mean zero and variance equal to the residual variance from the EM procedure.

### 2.4 Direct Estimation of Model Parameters

An alternative approach to handling missing data is to directly estimate the parameters of interest. For a fixed gene, in sample *j*, let *w_j_* denote an unobserved value of the gene expression *y_j_* and let *z_j_* be an indicator of an observed expression value as in Equation 1. The MLE of the variance for a given gene is:

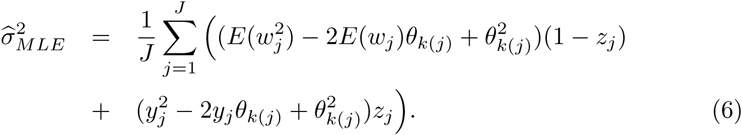

In contrast, the sample variance estimate following SI for a given gene is:

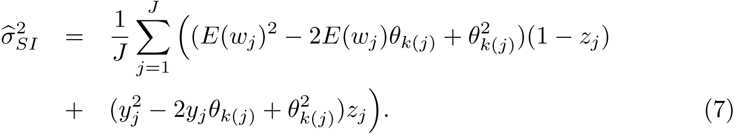

Derivations of these equations are given in Appendix A of the Supplementary Materials.

These equations differ exclusively in the first element within the summation: 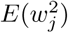 versus *E(wj*)^2^. Taking the difference between Equations (6) and (7) yields:

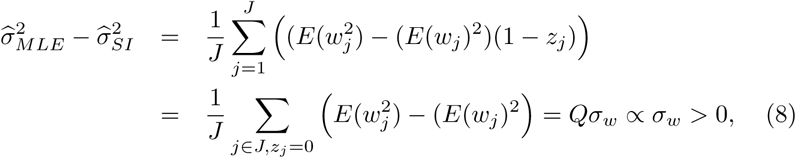

where *Q* > 0 is the proportion of non-detects for the given gene. Since 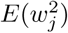 is greater then *E(w_j_*)^2^, the variance estimated after SI is smaller than the MLE of the variance. As the number of non-detects increases, Q increases, and hence the difference between 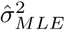 and 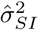 increases with the number of non-detected values. Note that these relationships hold for any given gene.

### 2.5 Simulation study design

We performed simulation studies with 16 or 90 genes, 6 sample-types, 4, 6, or 10 replicates within each sample-type, and a common missing data mechanism parameterized as in Equation 2. For each scenario we generated 100 complete data sets with the same gene expression *θ_ij(k)_*, within-replicate variance 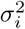, and parameters of missing mechanism (*β*_0_,*β*_1_). To summarize the performance of each estimation procedure, we report the 25^th^, 50^th^, and 75^th^ quantiles of each assessment measure for all genes and samples. For example, in the case of 16 genes and 6 sample-types, there are 16 different 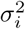 and 16x6=96 distinct values of *θ_ij(k)_* for each simulated data set.

## 3 Results

### 3.1 Simulation studies

#### 3.1.1 Single imputation underestimates the residual variance of gene expression

We expect the variability to be underestimated when performing a SI procedure. To confirm this and assess the bias and MSE of 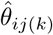 and 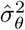, we performed a simulation study [4] with 16 genes as described in Section 2.5. While the SI bias and MSE are small for both *θ_ij(k)_* and 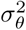, the SI bias for 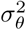 is always negative (Table 1). Note that for SI, 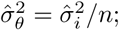 therefore, we confirm that the SI procedure underestimates the residual variance.

**Table 1.** Comparison of methods to handle missing data based on 100 simulated data sets.

|  | Bias |  |  | MSE |  |  |
| --- | --- | --- | --- | --- | --- | --- |
|  | Mean imputation |  |  |  |  |  |
| $\theta$ | -0.035 | -0.013 | 0.009 | 0.071 | 0.114 | 0.155 |
| $\sigma^2_\theta$ | -0.032 | -0.024 | -0.012 | 0.000 | 0.001 | 0.002 |
|  | Single imputation |  |  |  |  |  |
| $\theta$ | -0.024 | 0.003 | 0.024 | 0.072 | 0.114 | 0.156 |
| $\sigma^2_\theta$ | -0.014 | -0.007 | -0.004 | 0.000 | 0.001 | 0.001 |
|  | Multiple imputation |  |  |  |  |  |
| $\theta$ | -0.024 | 0.003 | 0.023 | 0.072 | 0.114 | 0.155 |
| $\sigma^2_\theta$ | -0.020 | -0.001 | 0.014 | 0.000 | 0.001 | 0.004 |
|  | Direct estimation |  |  |  |  |  |
| $\theta$ | -0.024 | 0.003 | 0.025 | 0.072 | 0.114 | 0.156 |
| $\sigma^2_\theta$ | -0.001 | 0.000 | 0.000 | 0.000 | 0.001 | 0.001 |

#### 3.1.2 Performance assessments of the proposed methods

To gain insights into the performance of the proposed MI and DirEst methodology, we constructed a simulation study (see Section 2.5) to compare the bias and MSE of the model parameter estimates from four imputation techniques: mean imputation, single imputation, multiple imputation, and direct estimation. Table 1 displays the 25^th^, 50^th^, and 75^th^ quantiles of the bias and MSE for 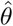 and 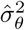 for all four methods. The mean imputation method underestimated both *θ* and 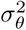 in this study. These disadvantages of mean imputation are noticeable even for the relatively small proportion of missing values in this study (6-11%). Single imputation performed almost as well as multiple imputation and direct estimation with respect to the bias and MSE of 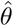; however, as expected SI underestimated 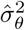. MI on average had similar performance to DirEst; however, the range of MI bias is generally wider than DirEst. In summary DirEst and MI are the most accurate methods in our assessments.

#### 3.1.3 Direct estimation is robust to misspecification of the missing data mechanism

We used the same simulated data to assess the robustness of our method to model misspecification. We compared the performance of the method under three possible link functions: logit, probit, and cloglog. Each simulation used the model described in Section 2.1. To assess the effect of the link function, we chose to fix the number of genes (*n*=16) and number of replicates (*k*=6) within each of the 6 sample types. In this case the correctly specified model is a logit link.

All three link functions yield very similar results in terms of bias and MSE for *θ* and *σ*^2^ (Table 2). Logit link gives estimates closer to the true values followed by cloglog then probit. Typically researchers are interested in estimating average gene expression *θ* and its variability *σ*^2^. All three link functions performed almost identically in estimating these parameters; therefore, the proposed method appears robust to the choice of link function in estimating gene expression means and variances in this simulation study. Because of the model robustness to the choice of the link function, in the remaining sections we present results using the logit link.

**Table 2.** Performance assessments of direct estimation under misspecification of the missing data mechanism based on 100 simulated data sets.

|  | Bias |  |  | MSE |  |  |
| --- | --- | --- | --- | --- | --- | --- |
| 16 genes | k=6 |  |  |  |  |  |
| logit |  |  |  |  |  |  |
| $\beta_0$ | -0.593 | | | 32.905 | | |
| $\beta_1$ | 0.014 | | | 0.027 | | |
| $\theta$ | -0.024 | 0.003 | 0.025 | 0.072 | 0.114 | 0.156 |
| $\sigma^2$ | -0.008 | -0.003 | 0.003 | 0.012 | 0.029 | 0.047 |
| probit |  |  |  |  |  |  |
| $\beta_0$ | 16.31 | | | 276.39 | | |
| $\beta_1$ | -0.459 | | | 0.219 | | |
| $\theta$ | -0.022 | 0.003 | 0.024 | 0.071 | 0.114 | 0.153 |
| $\sigma^2$ | -0.009 | -0.003 | 0.003 | 0.012 | 0.029 | 0.048 |
| cloglog |  |  |  |  |  |  |
| $\beta_0$ | 7.509 | | | 71.845 | | |
| $\beta_1$ | -0.228 | | | 0.065 | | |
| $\theta$ | -0.025 | 0.003 | 0.026 | 0.071 | 0.114 | 0.156 |
| $\sigma^2$ | -0.006 | -0.003 | 0.003 | 0.012 | 0.029 | 0.048 |

#### 3.1.4 The effect of the sample size

To assess the effect of sample size, we compared the performance on data sets with 16 or 90 genes and 4, 6, or 10 replicates for each sample type. An increase in the number of genes significantly improves the estimates of *β*_0_ and *β*_1_, while the accuracy of the parameters of interest, *θ* and *σ*^2^, remain basically the same (Table 3). These results are consistent across different numbers of replicates. Additionally, the MSE decreases as the number of replicates increase for both 16 and 90 genes. There is substantial bias and MSE in the estimates of **β*_0_* for 4, 6, and 10 replicates; however, these decrease from 16 to 90 genes. MSE decreases with an increase in the number of replicates, but stays consistently large. These high values of MSE for **β*_0_* have very little effect on the estimates of *θ* and *σ*^2^. Similar results were obtained for probit and cloglog links.

**Table 3.** Performance assessments of direct estimation for varying data set sizes based on 100 simulated data sets.

|  | 16 genes |  |  |  |  |  | 90 genes |  |  |  |  |  |
| --- | --- | --- | --- | --- | --- | --- | --- | --- | --- | --- | --- | --- |
|  | Bias |  |  | MSE |  |  | Bias |  |  | MSE |  |  |
|  | k=4 |  |  |  |  |  |  |  |  |  |  |  |
| $\beta_0$ | -2.101 | | | 78.176 | | | 1.018 | | | 7.895 | | |
| $\beta_1$ | 0.055 | | | 0.062 | | | -0.034 | | | 0.007 | | |
| $\theta$ | -0.025 | 0.006 | 0.033 | 0.111 | 0.173 | 0.226 | -0.022 | 0.009 | 0.045 | 0.121 | 0.172 | 0.253 |
| $\sigma^2$ | -0.012 | -0.008 | 0.011 | 0.020 | 0.047 | 0.091 | -0.018 | -0.004 | 0.009 | 0.025 | 0.045 | 0.098 |
|  | k=6 |  |  |  |  |  |  |  |  |  |  |  |
| $\beta_0$ | -0.593 | | | 32.905 | | | -0.107 | | | 5.427 | | |
| $\beta_1$ | 0.014 | | | 0.027 | | | -0.00018 | | | 0.004 | | |
| $\theta$ | -0.024 | 0.003 | 0.025 | 0.072 | 0.114 | 0.156 | -0.016 | 0.005 | 0.034 | 0.083 | 0.113 | 0.160 |
| $\sigma^2$ | -0.008 | -0.003 | 0.003 | 0.012 | 0.029 | 0.047 | -0.015 | -0.003 | 0.007 | 0.016 | 0.029 | 0.053 |
|  | k=10 |  |  |  |  |  |  |  |  |  |  |  |
| $\beta_0$ | -2.214 | | | 35.204 | | | -1.076 | | | 4.078 | | |
| $\beta_1$ | 0.060 | | | 0.028 | | | 0.029 | | | 0.003 | | |
| $\theta$ | -0.014 | 0.003 | 0.020 | 0.044 | 0.069 | 0.089 | -0.012 | 0.005 | 0.023 | 0.049 | 0.069 | 0.096 |
| $\sigma^2$ | -0.002 | 0.005 | 0.009 | 0.006 | 0.019 | 0.030 | -0.011 | -0.003 | 0.006 | 0.009 | 0.016 | 0.030 |

### 3.2 Comparison of proposed methods using real data

We applied the proposed methodology to three real datasets. The first dataset is composed of two cell types and three treatments (Sampson et al., 2013); the second dataset is a study of the effect of p53 and/or Ras mutations on gene expression (McMurray et al., 2008); the third dataset consists of nine gene perturbations with matched control samples (Almudevar et al., 2011). As in the original publications, all three datasets were normalized to a reference gene, Becn1. Additional details regarding each of these datasets can be found in the original publications.

#### 3.2.1 Difference in variance estimates between single imputation and direct estimation

We compared estimates of the variance from a direct estimation procedure with SI estimates for absolute and relative quantification. The minimum, maximum, first, second and third quartiles for 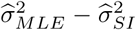are presented in Table 4 for all three datasets. The difference between the two variance estimates is usually small, but in Dataset 2 the variance estimates for the gene *Afp* differ by 35.38. In this dataset *Afp* has 13 non-detects out of 14 values. This is a concrete example of the difference in variance estimates increasing as the number of non-detects increases, in other words, the effect of a larger *Q* in Equation 8.

**Table 4.** Summary statistics for the difference between estimates of within replicate variance: 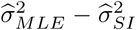 in three real data sets.

|  | Min. | 1<sup>st</sup>.Qu. | Median | Mean | 3<sup>rd</sup>.Qu. | Max. |
| --- | --- | --- | --- | --- | --- | --- |
| Dataset 1 | 0.024 | 0.026 | 0.033 | 0.095 | 0.109 | 0.429 |
| Dataset 2 | 0.006 | 0.009 | 0.027 | 1.014 | 0.110 | 35.380 |
| Dataset 3 | 0.010 | 0.019 | 0.040 | 0.056 | 0.042 | 0.199 |

The individual differences in variance estimates for absolute and relative expression can be seen in Supplementary Figures 1 and 2 in Appendix C of the Supplementary Materials, respectively. Similar to the results show in Table 4, most of the differences are fairly small with a few exceptions, especially in Dataset 2. However, even small differences may influence downstream analyses and lead to anti-conservative inference.

**Figure 1:**
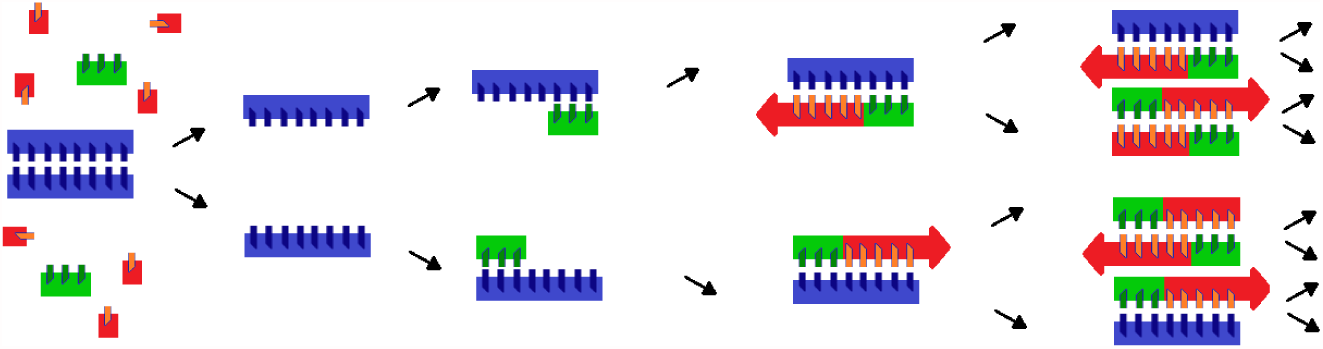
Polymerase chain reaction begins with a mixture of double-stranded DNA (blue), DNA primers (green), and nucleotides (red). The reaction follows three main steps: (1) denaturation: heating disrupts the bonds between DNA nucleotides to produce single-stranded DNA molecules, (2) annealing: DNA primers bind to their complementary single-stranded DNA sequence and begin DNA formation, (3) elogation: DNA polymerase synthesizes a new DNA strand complementary to each single-stranded DNA sequence. This sequence of steps, called a *cycle,* results in a doubling of the initial DNA molecules under ideal conditions.

**Figure 2:**
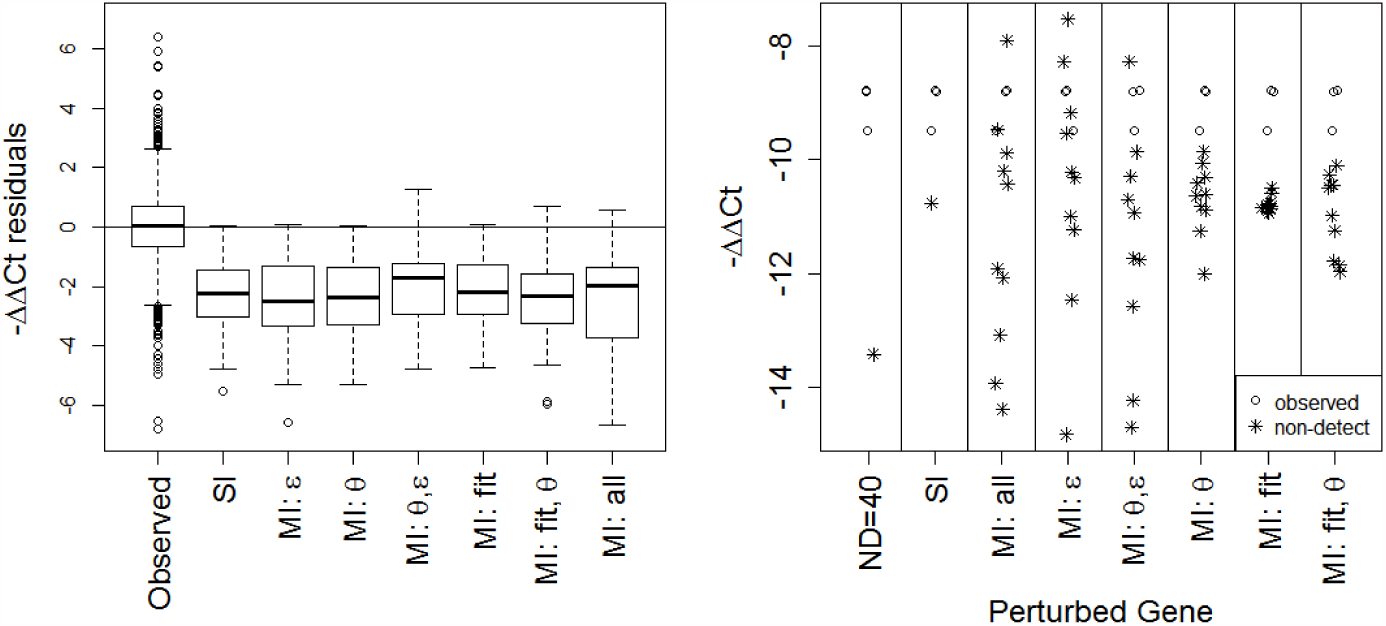
Comparison of single and multiple imputation using a real data example. The left panel shows within-replicate residuals stratified by the presence and handling of non-detects. The average ∆∆Ct values were calculated over the replicates for gene i and sample j. The residuals for each gene and sample-type were summarized and are plotted here. The box-plots from left to right display the distribution of residuals for: the observed data, missing data after SI, and missing data after MI with different sources of variability. The right panel shows the responses of gene Sema7a to the perturbation of Wnt9a from Dataset 3. Imputed values are denoted with an asterisk. ∆∆Ct values produced by replacing a non-detect with a value of 40 are in the leftmost panel, SI estimates are in the next panel, and estimates resulting from applying MI with different combinations of variability sources are shown in the rightmost six panels.

#### 3.2.2 Multiple imputation for absolute and relative quantification

We have implemented a MI procedure in the presence of non-detects for absolute quantification. To account for the uncertainty in the imputed values, we propose to use different variability sources and combinations thereof. The MI approach allows one to choose the number of complete data sets to impute for a set of non-detects, in contrast to SI which returns one complete data set of observed and imputed values. In Supplementary Figure 3 in Appendix C of the Supplementary Materials, we show gene expression estimates produced by SI and MI for Datasets 1 and 2. MI can incorporate all sources of uncertainty at once (“MI: all”), pairs of sources (“MI: *θ*, *ε*”, “MI: fit, *θ*”) or one source at a time (“MI: *ε*”, “MI: *θ*”, “MI: fit”).

In Supplementary Figure 4 in Appendix C of the Supplementary Materials, we show the distribution of residuals for the observed data and the results of a SI procedure and several MI procedures for the missing data. Because a missing value likely represents slightly lower gene expression compared to the observed values from replicate samples, the majority of the residuals for the imputed values are slightly negative. While the medians are similar between the MI and SI results in both Datasets 1 and 2, the MI residuals often have a larger IQR because they incorporate additional sources of variability that are ignored by the SI procedure. The smallest impact on the distribution of the residuals is the uncertainty in the missing data mechanism, indicated as “fit” in Supplementary Figure 4, followed by uncertainty in *θ*. The biggest impact is measurement error, *ε*. Overall, the MI procedure better captures the uncertainty in estimates of absolute quantification.

Similar to absolute quantification, we have implemented an MI procedure in the presence of non-detects for relative quantification. While Datasets 1 and 2 focused on absolute expression, Dataset 3 focused on relative expression. The results for ten imputed data sets are presented in Figure 2. The distribution of within replicate residuals in MI compared to SI appears to be very similar in the case where only uncertainty in *θ* is included in MI. Overall, the mean of the residual distribution stays relatively unchanged, but the IQR is wider due to the incorporated sources of variability in the model. In summary, the uncertainty in estimates of relative quantification are better captured by MI than by SI.

## 4 Discussion

This paper has introduced two methods to account for missing data in qPCR experiments: multiple imputation and direct estimation. Both methods treat qPCR non-detects as missing not at random, and model the missing data mechanism with a two parameter sigmoidal curve. Using simulations, we showed that DirEst and MI can accurately estimate the first two moments of the distribution of gene expression and out perform SI and mean imputation. Additionally, we showed that the proposed methods are robust to model misspecification and perform fairly well even with small sample sizes and without a large number of replicate samples.

Expanding upon a previous SI procedure for non-detects, we developed a MI procedure that incorporates different sources of uncertainty into the model. This approach is preferred when the actual expression estimates are required for analysis, for example in gene regulatory network modeling, clustering, or coexpression analysis. We also developed a method to estimate model parameters directly, DirEst approach. This method can be used when the mean and variance are sufficient statistics, e.g. when performing analysis of differences in average gene expression across groups.

A previous SI procedure, described in [18], preserves the first moment but introduces bias in the estimation of the variance. We have shown that this underestimation of the variability increases with the proportion of non-detects. Smaller estimated standard deviations lead to smaller p-values, which result in anti-conservative inference and a larger Type I error. This can lead to reporting significant results where there is no statistical difference.

Future work should further investigate these approaches and extensions of these methods. One current limitation is that the proposed methods require an observed value for a given gene in at least one replicate sample; however, it is possible for all the replicates of a given sample-type to be non-detects. Furthermore, it is important to distinguish between the lack of gene expression and the lack of detection of an expressed gene. Additionally, in this work we assume that the gene expressions are non-correlated; posing a correlation structure would allow us to borrow information across genes and improve parameter estimation. Finally, the use of qPCR data in medical applications requires methods that do not rely on replicate samples.

In conclusion, both MI and DirEst models are presented as strong options for data with non-detects in qPCR when repeating the experiment is not feasible. By using them we may be able to make even more robust inference in downstream analysis and have more confidence in generating scientific hypotheses.

## 5 Software

Our algorithm is implemented in the R/Bioconductor package, *nondetects*. Our software can handle a variety of study designs. For example, if samples are collected in batches and there is a control sample for each batch, the software can adjust for batch effects. To clearly describe the functionality of the package and demonstrate multiple reproducible real data examples, we created a user friendly vignette to accompany the package. Software in the form of R code for the simulation studies and real data applications, together with input data sets, is available in Appendix D of the Supplementary Materials.

## 6 Supplementary Material

Supplementary materials contain derivations of the variance estimates for SI and MLE and their difference in Appendix A. In Appendix B we present tables with the simulation results; Supplementary Figures are shown in Appendix C.

## 7 Acknowledgments

The authors thank Ollivier Hyrien and Andrew McDavid for comments and feedback on the earlier versions of this manuscript and for additional advice. *Conflict of Interest:* None declared.

